# Identifying Critical Points of Trajectories of Depressive Symptoms From Childhood to Young Adulthood: Evidence of Sex Differences from the Avon Longitudinal Study of Parents and Children (ALSPAC)

**DOI:** 10.1101/193680

**Authors:** Alex S. F. Kwong, David Manley, Nicholas J. Timpson, Rebecca M. Pearson, Jon Heron, Hannah Sallis, Evie Stergiakouli, Oliver S.P. Davis, George Leckie

## Abstract

Depression is a common mental illness associated with increased substance misuse and risk of suicide. Potential risk factors for depression include sex and depressive symptoms in early life, however the mechanisms responsible are not yet understood. Research has focused on late childhood and adolescence as this developmental period may be a modifiable risk factor that prevents or reduces depression at a later stage. It is also important to establish at what ages the level of depression is changing as this will help identify critical points to intervene with treatment. We used multilevel growth-curve models to explore adolescent trajectories of depressive symptoms in the Avon Longitudinal Study of Parents and Children, a UK based pregnancy cohort. Using data from 9301 individuals, trajectories of depressive symptoms were constructed for males and females between 10.6 and 22.8 years old. We calculated the age of peak velocity for depressive symptoms (the age at which depressive symptoms increases most rapidly) and the age of maximum depressive symptoms. Adjusted results suggested that being female was associated with a steeper trajectory compared to being male (per 1 year increase in relation to depressive symptoms: 0.128, SE = 0.035, [95% CI: 0.059, 0.198]; *p* <0.001). We found evidence suggesting that females had an earlier age of peak velocity of depressive symptoms (females 13.7 years old, SE = 0.321, [95% CI: 12.9, 14.4] and males 16.4 years old, SE = 0.096, [95% CI: 16.2, 16.6]; *p* <0.001), but weak evidence of an earlier age of maximum depressive symptoms (*p* = 0.125). Possible mechanisms that underlie this sex difference include the roles of pubertal development and timing. Using multilevel growth curve models to estimate the age of peak velocity and maximum depressive symptoms for different population subgroups may provide useful knowledge for treating and preventing later depression.

## Introduction

Depression is a common mental illness which is believed to affect more than 300 million people worldwide [1]. Depression is comorbid with other psychiatric conditions including anxiety, bipolar disorder and schizophrenia, and often associated with increased substance misuse, impaired educational attainment and increased risk of suicide [2-6]. Identifying the mechanisms underpinning depression has become increasingly important for identifying treatments and interventions.

One avenue for research has focused on childhood and adolescent depression as a potentially modifiable risk factor for adulthood depression [7-9]. Evidence suggests this developmental period of late childhood and adolescence is important for subsequent mental health and identifying and treating depression during this time may limit or prevent depression in later life [2, 9, 10]. However, the critical age to intervene in order to avoid later depression had not yet been established. Assessing depression in childhood and adolescence can also be difficult due to comorbidity and the suggestion that depression exists on a continuum showing varying levels of subclinical and clinical depression [4, 11-13]. Continuous measures of depressive symptoms can be easier to collect and can predict clinical depression at a later stage [9, 14]. In this context, identifying depressive symptoms early on in development is useful for characterising population-based patterns of emergence and for early identification of at risk individuals [2, 15].

Longitudinal studies can be used to observe how the level of depressive symptoms changes over time [4, 7, 10]. However, some studies only use cross-sectional data or data with few time points, which makes it hard to understand the nature of change [16]. To appropriately examine changes in depressive symptoms, longitudinal studies should use data from multiple timepoints across a wide enough time period. Trajectories or growth curve models can then be used to study how depressive symptoms change across time and how such patterns of change vary across individuals. These trajectories contain pertinent features such as point of maximum growth and peak velocity (i.e., the point in which a trait is increasing most rapidly) [17, 18]. It is also possible to associate these trajectories with earlier exposures [19, 20] or later distal outcomes [16, 21-23]. Trajectories can capture the complex variability of change over time and delineate these trajectories to examine individual and population growth patterns [7, 11, 12, 24-26], thus making them useful tools for understanding how depressive symptoms may change. However, no studies have specifically examined the age of peak velocity of depressive symptoms (i.e., the age at which depressive symptoms are increasing most rapidly) and how this might vary across population subgroups. By calculating age of peak velocity, it may be possible to infer how depressive symptoms change over time across the population, and then identify at-risk individuals and implement early stage interventions.

Existing research using trajectories has demonstrated that features including age of peak depressive symptoms could be calculated, but in most cases, this would come from graphs or via latent class growth modelling (LCGM) [27] or general growth mixture modelling (GGMM) [28, 29]. The limitation with these approaches is that one is left comparing multiple subpopulation trajectories (often between 3 and 5) which may not be translatable. Additionally, whilst the LCGM and GGMM trajectories are useful for explaining changes in depressive symptoms, stratification into latent subpopulations may be problematic when assumptions about the data are not met or with small sample sizes [30]. These methods can also create subgroups that do not exist and in certain conditions may be inapplicable for traits that constantly fluctuate [30]. One alternative approach is to use multilevel growth-curve models to predict individual specific trajectories [31, 32]. This approach has been widely applied to measure other forms of childhood growth, for example height and weight [19, 21]. Issues related to sample sizes and fluctuations in the data are circumvented with multilevel growth-curve models. Moreover, multilevel growth-curve models provide trajectories for specific populations which capture individual variation and could help researchers explore how depressive symptoms change over time and how variable depressive symptoms may be.

Existing literature suggests that depressive symptoms trajectories are heterogenous with multiple trajectories [7, 11, 12, 25, 33, 34]. These trajectories can be affected by multiple risk factors including early stressful life events [35], pubertal timing [36], parental relationships [37] and social economic status [7]. However, one of the most potent covariables for depressive symptoms is sex [2, 25, 38, 39]. Multiple studies suggest fundamentally different trajectories exist between males and females, and that females have steeper trajectories, and commence on these trajectories earlier than males [7, 12, 25, 33]. Natsuaki et al., [36] used multilevel growth curve modelling to examine sex differences in depressive symptoms and observed that females had steeper trajectories and commenced on these trajectories earlier than males. They also observed a peak of depressive symptoms towards late adolescence that was steeper for females. Studies using LCGM or GGMM have found somewhat similar and comparable results. Chaiton et al., [12] found three distinct trajectories which they labelled ‘low’, ‘moderate’ and ‘high’ for both males and females, whilst Dekker et al., [25] found six trajectories which differed slightly for males and females ranging from what they term ‘very low decreasing’ to ‘high increasing’ with other trajectories indicating both a ‘peak in childhood’ and ‘peak in adolescence’. Fernandez Castelao…Kroner-Herwig [33] also found four trajectories for females and three for males, each varying from ‘low to increasing’ to ‘high to decreasing’. The evidence suggests that females have steeper trajectories, yet the reasons behind this are still not established and more longitudinal research is needed to explain these sex differences.

Understanding the nature of how depressive symptoms change through the use of multilevel growth-curve models may be one method for further explaining these differences. However it is difficult to interpret the nature of trajectories of depressive symptoms or make inferences about future depression in many studies where data are not collected past a certain age (e.g., beyond 16 years where the data may be increasing, but no longer measured). It is also challenging to understand elements of the trajectory, or whether the depressive symptoms are fluctuating if only a snapshot of the period of growth is measured. To fully understand the nature of depressive symptoms trajectories and how they change, it is important to examine a wealth of data across a wide enough time period (e.g., from childhood to adulthood). Some progress has been made with the data available, yet most of these studies have used cross-sectional methods to infer how depressive symptoms change over time. Longitudinal research focusing on the transition of depressive symptoms between childhood, through adolescence and into adulthood will aid in our understanding of how depressive symptoms change, and how we can use this information for treatment and prevention.

In the current study, we aimed to build upon this literature and use multilevel growth-curve models to estimate sex-specific trajectories of depressive symptoms in a UK-based population cohort with data ranging from late childhood (approximately 10 years old) to young adulthood (approximately 23 years old). The primary objective was to examine how trajectories differ between males and females. The second aim was to ascertain the age of peak velocity for depressive symptoms and age of maximum depressive symptoms and whether these differ between males and females. Finally, we aimed to estimate average depressive symptoms scores at the age of peak velocity and maximum depressive symptoms in order to better understand the severity of depression at these ages.

## Methods

### Participants

Participants were from the Avon Longitudinal Study of Parents and Children (ALSPAC), an ongoing population-based study in the South-West of England designed to examine the effects of a number of factors on health and development. ALSPAC recruited pregnant mothers with an estimated delivery date between April 1991 and December 1992. The initial cohort consisted of 14,541 pregnancies, with 13,988 children still alive one year later (52% males and 48% females). ALSPAC children were slightly more educated at age 16 compared to the national average, slightly more likely to come from homeowner backgrounds and slightly more likely to be of Caucasian descent [40]. Ethical oversight for the study was obtained from the ALSPAC Law and Ethics Committee and Local Research Ethics Committees. Participant data has been collected on the mothers, fathers and children from early pregnancy, and measured via questionnaires and regular clinic visits. Part of this data was collected using REDCap (https://projectredcap.org/resources/citations/). Detailed information about ALSPAC is available on the study website (www.bris.ac.uk/alspac), which also contains a fully searchable data dictionary (www.bris.ac.uk/alspac/researchers/data-access/data-dictionary).

### Measures

#### Depressive symptoms

The outcome measure of the trajectories were depressive symptoms, measured using the short mood and feelings questionnaire (SMFQ) [41]. The SMFQ is a 13-item questionnaire that measures the occurrence of depressive symptoms over the preceding two weeks with higher scores indicating more severe depressive symptoms (range 0-26). The SMFQ correlates highly with clinical measures of depression [4], and has been used as a viable measure of depressive symptoms in previous epidemiological studies [14, 41]. We used self-report SMFQ data from eight occasions beginning in late childhood and extending to early adulthood. Questionnaires were either completed via postal questionnaire or by computer at a clinic visit. The sample size at each measurement occasion decreased from 7735 at 10.6 years to 3353 at 22.8 years (Table 1). See table 1 for SMFQ descriptive statistics.

**Table 1.**
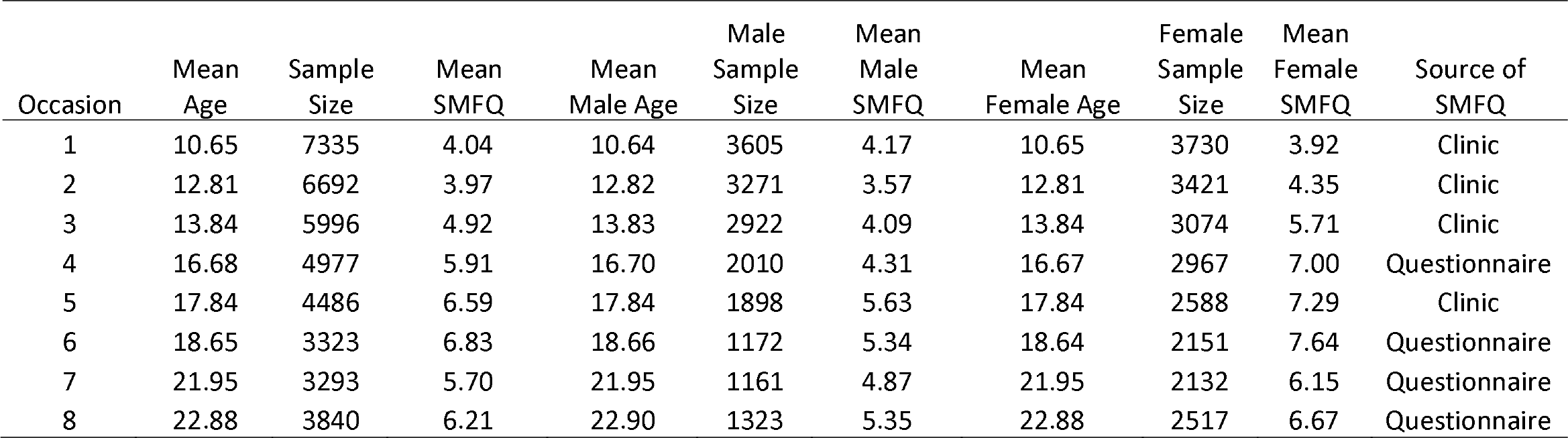
Descriptive statistics of the Short Mood and Feelings Questionnaire (SMFQ).

#### Exploratory variables and covariables

Our main exploratory variable was sex, identified from birth notifications around the time of delivery and coded as a dummy variable for being female. Participants were removed from the analysis if sex was unknown. Confounders were included based upon previous evidence from the depressive symptoms literature that highlight correlations between demographic variables with depressive symptoms. These confounders were completed by the participant’s main carer and assessed during the antenatal period. These included: maternal education (coded as ‘A-level or higher’, ‘O-level’ or ‘<O-level’), maternal social class (coded as ‘Professional occupations or managerial and technical occupations’ or ‘Skilled non-manual occupations, skilled manual occupations, partly-skilled occupations and unskilled occupations’, parity (whether the study child was 1^st^/2^nd^/3^rd^ born or greater), housing tenure (coded as ‘Mortgaged or owned’, ‘Privately rented’ or ‘Subsided rented’), financial difficulties (yes/no), maternal smoking in pregnancy (yes/no), maternal prenatal depression (yes/no) and maternal postnatal depression (yes/no). A binary indicator of the SMFQ source was also included as a confounder (clinic/questionnaire).

### Statistical Methods

We estimated trajectories of depressive symptoms using a multilevel cubic polynomial growth-curve model. The constant, linear, quadratic and cubic terms are all allowed to vary randomly across individuals. The model specifies separate population-averaged growth curves for males and females. The model can then be written as

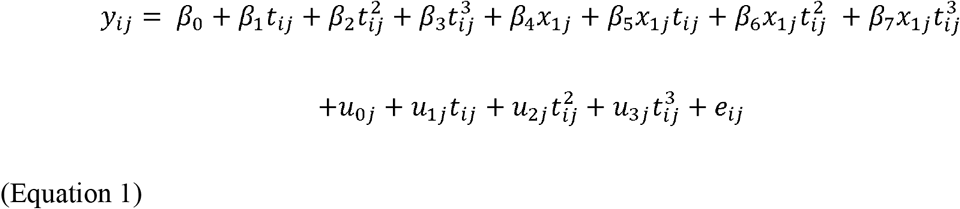

where *y_ij_* is the depressive symptom score and *t_ij_ y_ij_* is the age (centred around 16 years, the approximate sample mean) for individual *j* at occasion *i*. *x_ij_* is a dummy variable for being female, and *u_0j_*, *u_1j_*, *u_2j_*, and *u_3j_* are the random linear, quadratic and cubic effects, respectively. The occasion-specific residual *e_ij_* allows the depressive symptom scores to deviate from the perfectly cubic trajectories.

The random effects are assumed multivariate normal distributed with zero mean vector and constant covariance matrix

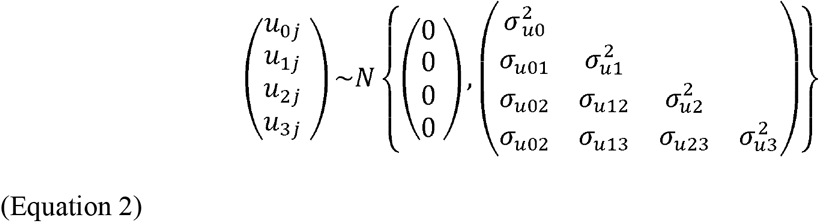

The residuals are assumed normally distributed with zero mean and sex specific variance

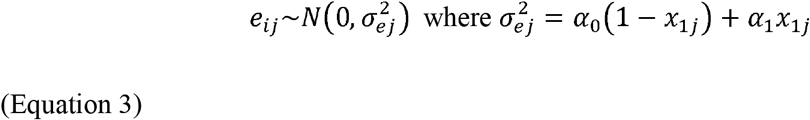

All analyses were conducted using Stata 13 (StataCorp, College Station, TX, USA) using the user-written runmlwin command [42], which calls the standalone multilevel modelling package MLwiN v2.35 (www.cmm.bristol.ac.uk/MLwiN/index.shtml). Stata code is provided in the supplementary materials.

#### Trajectory features

The population male and female trajectories are therefore given by

Male: 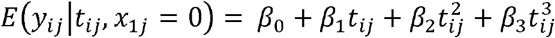

Female: 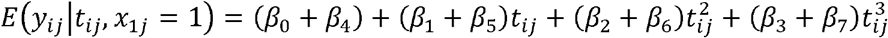

Several features were calculated from the mean trajectories including: the age of peak velocity of depressive symptoms (i.e., age at which depressive symptoms is increasing most rapidly), and the age of maximum depressive symptoms (i.e., age at where depressive symptoms were the highest on the trajectory). The depressive symptoms scores were also calculated at the age of peak velocity and the age of maximum depressive symptoms. These were calculated by differentiating equation 1 with respect to age and then calculated separately for males and females, and compared using the delta method in Stata (Supplementary Information).

## Results

### Sample description and characteristics

There were 9301 individuals with data on sex and at least one measurement of the SMFQ, resulting in 39,942 measurements (17,362 male/22,580 female). The total sample that included all the confounders was 6097 individuals, resulting in 27,952 measurements (12,362 male/15,590 female).

### Model selection

Descriptive statistics indicated the change in depressive symptoms followed a non-linear pattern (Table 1). Thus we needed to model age in a way that appropriately captured the nonlinearity of these trajectories. We chose a cubic polynomial model for two reasons: the first is the cubic term fitted the data better and seemed more plausible than the quadratic and quartic functions of age (Supplementary Information). The second reason is that compared to more flexible cubic splines and fractional polynomials which have also been used to model nonlinear growth [21], the cubic polynomial model is more parsimonious, especially when calculating trajectory features (Figure 1).

**Figure 1.**
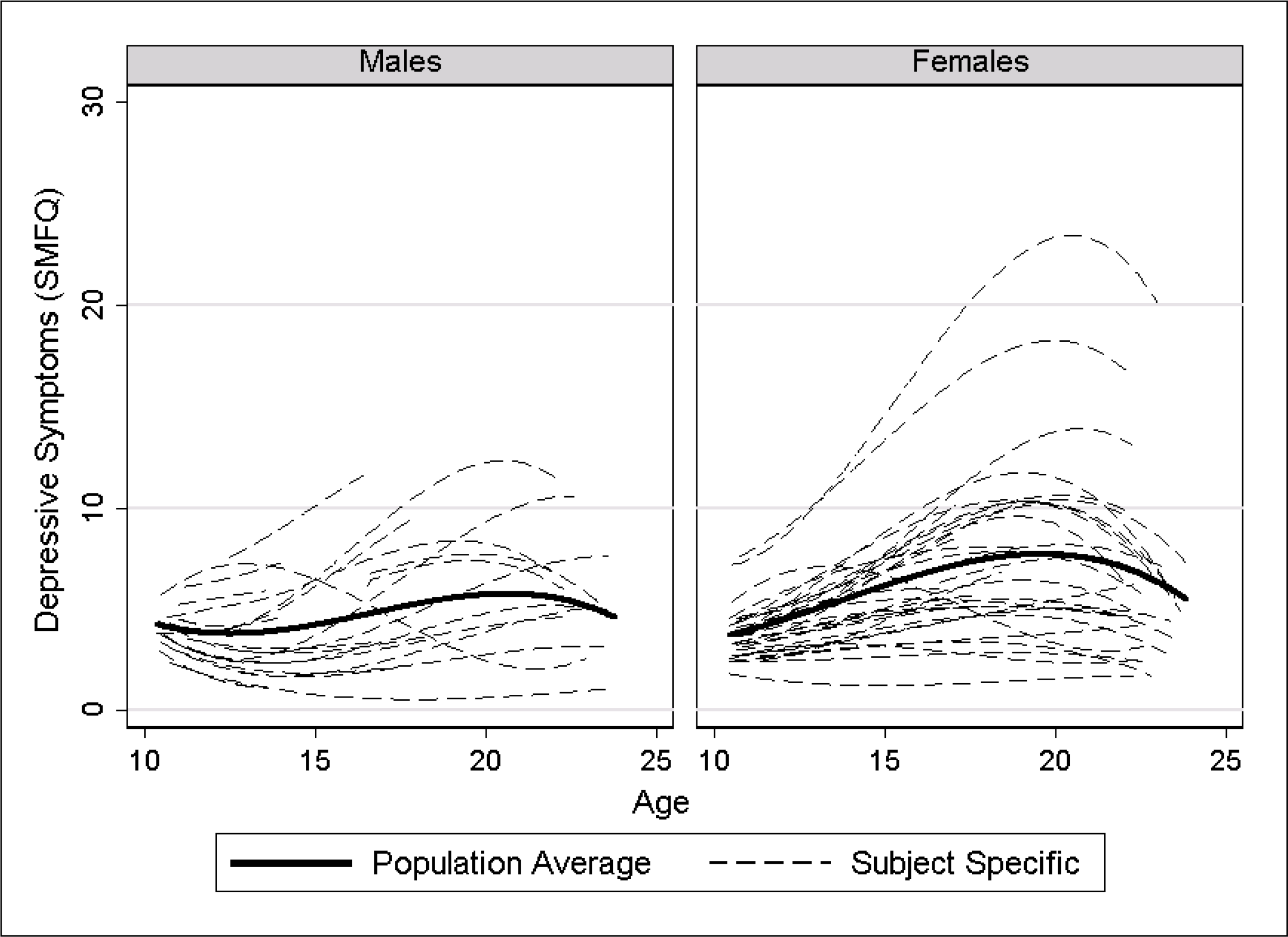
Individual and population trajectories for a random set of 100 participants. SMFQ: Short Mood and Feelings Questionnaire.

### Trajectories of depressive symptoms

Females and males followed distinctively different trajectories (Figure 1). The intraclass correlation evaluated at age 16 was 0.56 (SE = 0.008) suggesting substantial interindividual variation in depressive symptom scores and the need for multilevel growth-curve modelling. The correlation between the intercept (i.e., first measurement of depressive symptoms) and the slope (i.e., change in depressive symptoms with every year increase) was 0.52 (SE = 0.025) suggesting that individuals with higher depressive symptoms at age 16 had were also at that age experiencing more rapid increase in depressive symptoms. Table 2 shows the regression coefficients from the unadjusted model, along with the addition of confounders for the adjusted model (Supplementary Table 2). The Trajectories did not differ substantially with the addition of the confounders (Figure 2), so only the adjusted trajectories and adjusted main features of the trajectories are described from here onwards.

**Figure 2.**
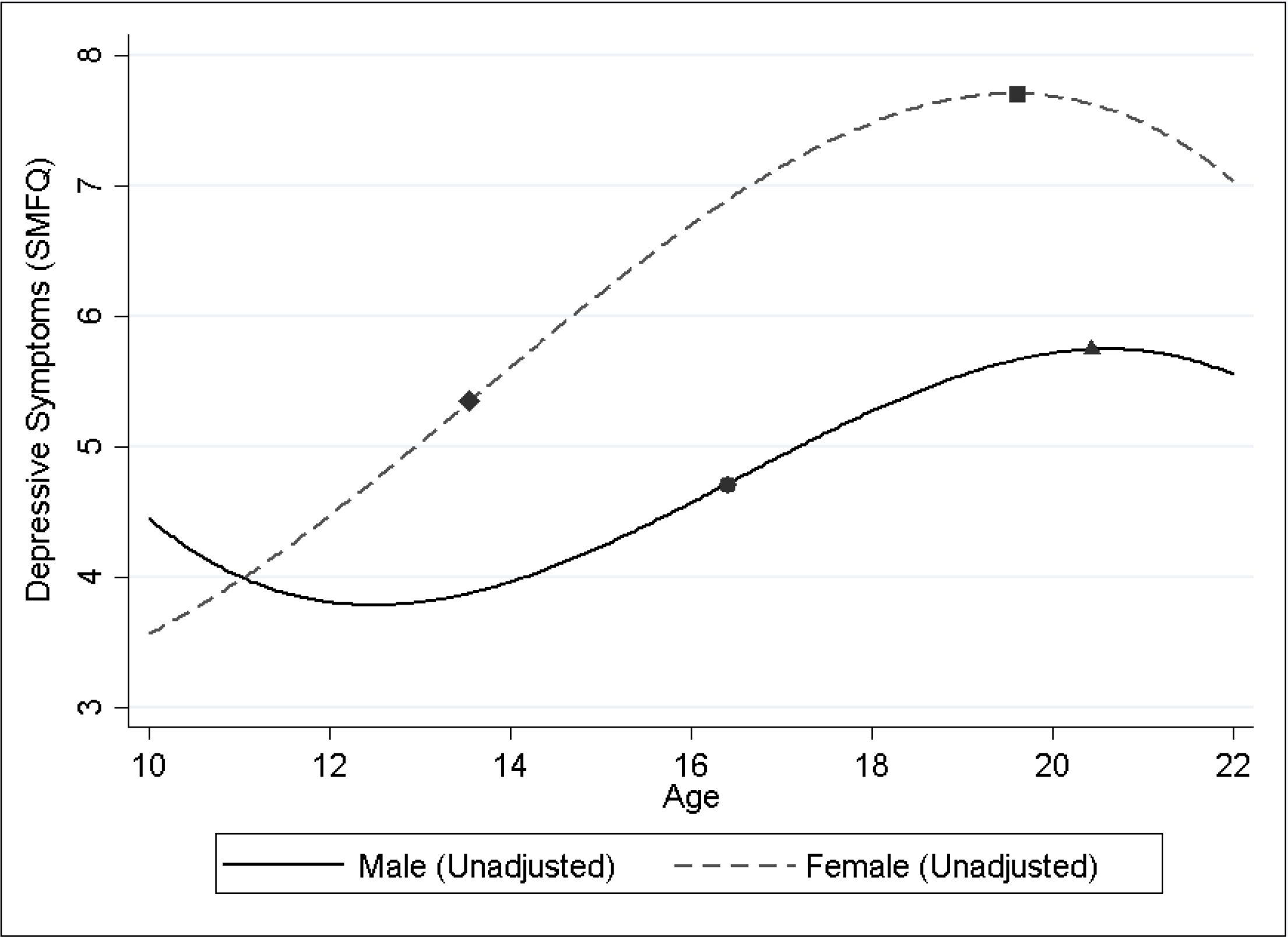
Unadjusted population trajectories for males and females. SMFQ: Short Mood and Feelings Questionnaire. Features of the trajectories are overlaid with the following terms:…Male age of peak velocity of depressive symptoms. ▴ Male age of maximum depressive symptoms. ♦ Female age of peak velocity of depressive symptoms. ▪ Female age of maximum depressive symptoms.

**Figure 3.**
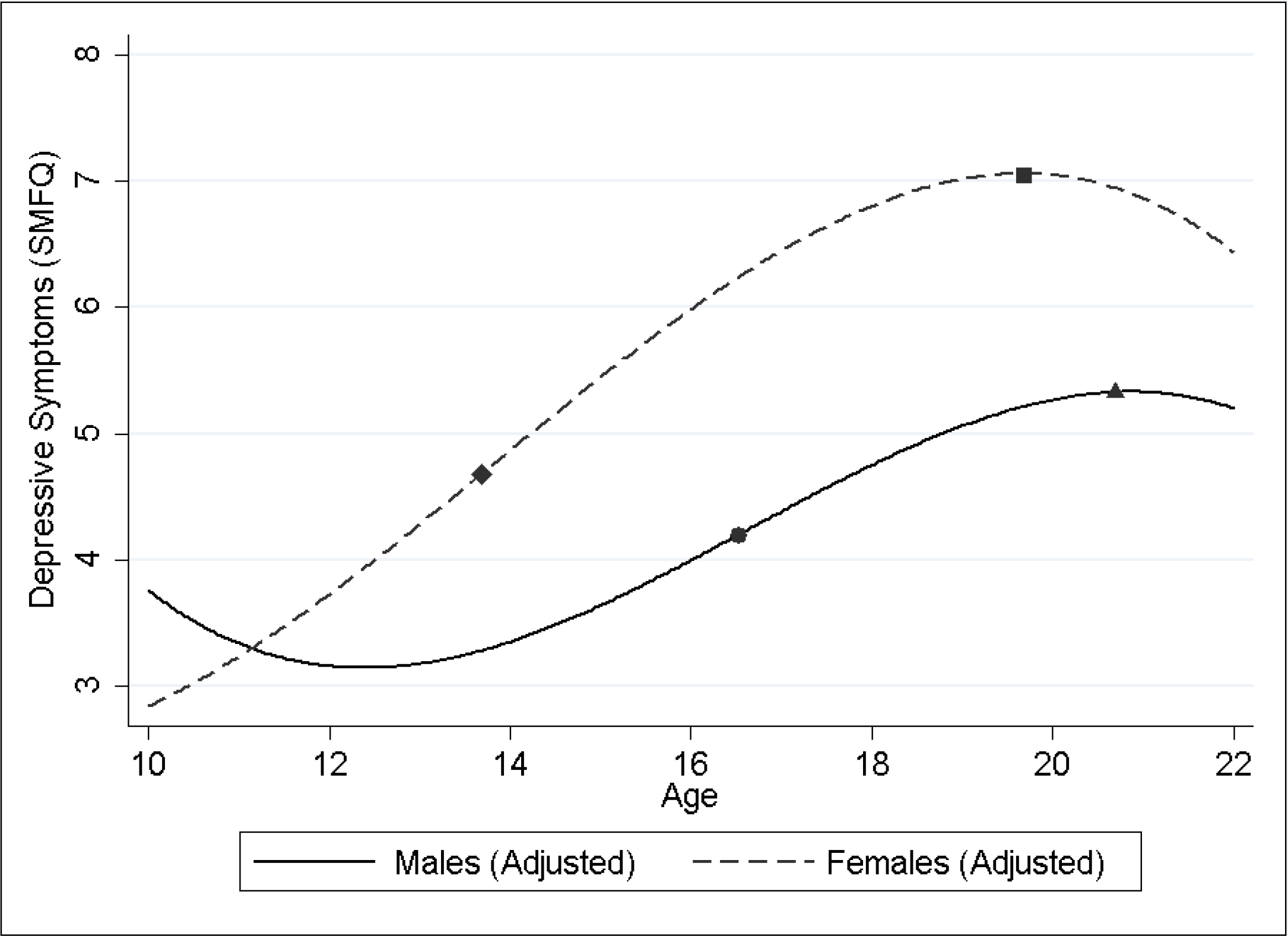
Adjusted population trajectories for males and females. SMFQ: Short Mood and Feelings Questionnaire. Features of the trajectories are overlaid with the following terms:…Male age of peak velocity of depressive symptoms. ▴ Male age of maximum depressive symptoms. ♦ Female age of peak velocity of depressive symptoms. ▪ Female age of maximum depressive symptoms.

Females were associated with higher depressive symptoms at 16 years of age (*β_0_* + *β_4_* = 5.987, SE = 0.11 [95% CI: 3.776, 4.218]) compared to males (*β_0_* = 3.997, SE = 0.11 [95% CI: 5.771, 6.202]; *p*^diff^ < 0.001). Females also had steeper trajectories compared to males (per 1 year increase in relation to depressive symptoms difference: 0.128, SE = 0.036 [95% CI: 0.059, 0.198]; *p*^diff^<0.001), except between 10 and 11 years old where males were predicted to have higher depressive symptoms. The female depressive symptoms trajectory was characterised by a strong increase throughout late childhood and into adolescence, followed by a peak in late adolescence. The female trajectory then continued to decrease into young adulthood. The male depressive symptoms trajectory started higher than the females, but decreased in late childhood before rebounding with a steady increase in early adolescence. The male trajectory continued to rise until a peak in early adulthood, which then followed a decline into young adulthood. There was evidence of Interactions between sex and the intercept and slopes respectively (Table 2)

**Table 2.**
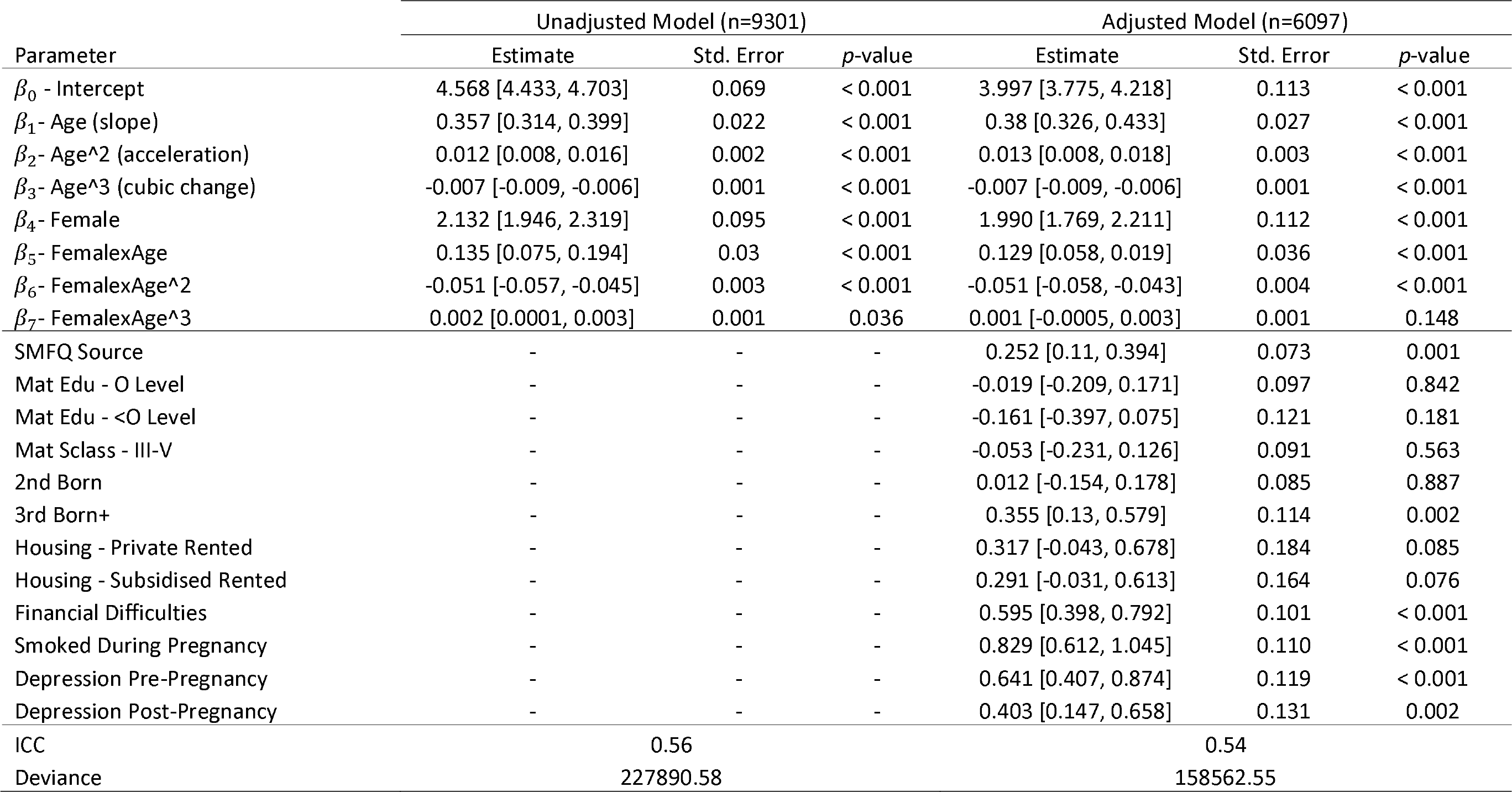
Regression coefficients for the unadjusted and adjusted models. 95% confidence intervals given in [parenthesis].

### Age of peak velocity and age at maximum depressive symptoms

The age of peak velocity in the adjusted model was earlier for females (13.66 years old, SE = .36 [95% CI: 12.97, 14.36]) compared to males (16.44 years old, SE = .12 [95% CI: 16.21, 16.67]; *p* <.001) (Table 3). However there was weak evidence of a difference between the age at maximum depressive symptoms for females (19.68 years old, SE = .58 [95% CI: 18.54, 20.82]) compared to males (20.68 years old, SE = .18 [95% CI: 20.32, 21.02]; *p* < .125).

### Depressive symptoms score at age of peak velocity and at maximum point of depressive symptoms

The predicted depressive symptoms scores at the estimated age of peak velocity were higher for females (4.77 points, SE = .12 [95% CI: 4.54, 5.0]) than they were for males (4.24 points, SE = .11 [95% CI: 4.01, 4.45]; *p* < .001) (Table 3). The depressive symptoms scores at the maximum point depressive symptoms were also higher for females (7.06 points, SE = .12 [95% CI: 6.81, 7.3]) compared to males (5.33 points, SE = .13 [95% CI: 5.07, 5.59]; *p* < .001).

**Table 3.**
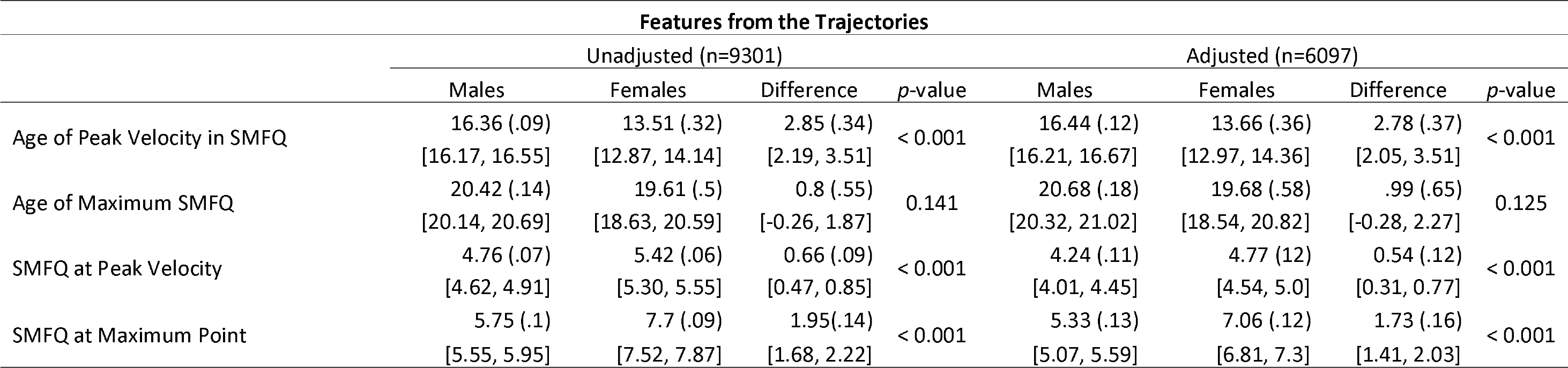
Calculated features from the trajectories for the unadjusted and adjusted models. Standard errors are given in (parenthesis), 95% confidence intervals are given in [parenthesis].

### Sensitivity analysis

Our initial analysis included individuals with at least one measurement from the SMFQ. Although individuals with only one measurement do contribute to the analysis (e.g. via association between depressive symptoms and sex), it is not meaningful to predict and examine trajectories for individuals with only one measurement. Therefore we ran a sensitivity analysis where we refitted our models on the subset of individuals who had three or more depressive symptom scores. The unadjusted sample size reduced from 9301 to 6878, and the adjusted from 6097 to 4800. However, removing these individuals had no substantial impact on our trajectories or the interpretation and conclusions from these trajectories (Supplementary tables 3 and 4).

## Discussion

The results indicate that females and males had distinctively different mean trajectories of depressive symptoms between late childhood and young adulthood. More specifically, females were associated with steeper trajectories of depressive symptoms compared to males, with the exception of between 10 and 11 years old where males had higher depressive symptoms. We also found strong evidence that females had an earlier age of peak velocity of depressive symptoms (i.e., age at which depressive symptoms is increasing most rapidly), and weak evidence that females had an earlier age of maximum depressive symptoms (i.e., age at which depressive symptoms is highest on the trajectory) in comparison to males. Finally we found that the depressive symptoms scores at the age of peak velocity and age of maximum depressive symptoms were higher for females compared to males, with distinct differences at both occasions.

Our findings support and extend previous research on the role of sex on trajectories of depressive symptoms throughout childhood to adulthood. For instance, steeper trajectories of depressive symptoms in females have been observed in multiple studies [7, 16, 33], and being female appears to be a considerable risk factor for depression [2, 39]. Our results support this as females were associated with steeper trajectories of depressive symptoms from age 11 onwards. Additionally, females were also associated with higher depressive symptoms scores at both the age of peak velocity and age of maximum depressive symptoms. Whilst this is unsurprising, it is important to corroborate and extend this research with findings that use longitudinal data across a wide time period and not just adolescence, for example. This will allow researchers and clinicians to better understand the nature of depressive symptoms and how they change over time. To date, only a handful of studies have had the capacity to use such methods [11, 24, 36] and similar findings are observed, resulting in a consistent pattern of results that highlight that females are at a higher risk of depression than males.

By using rich longitudinal data, we were also able to expand upon previous research by calculating the age of peak velocity and age of maximum depressive symptoms, as well as the depressive symptoms scores at both of these ages. Our results show that the age of peak velocity of depressive symptoms was almost 3 years earlier for females and calculating this age has the potential to be a useful tool for identifying at risk individuals and potential treatment. We infer that depressive symptoms are increasing most rapidly for females at 13.7 years old and for males at 16.4 years old. This suggests the potential to implement treatment and interventions at different ages for males and females to prevent these increases from continuing into later adulthood depressive symptoms or clinical depression. It may be clinically useful to identify features such as the age of peak velocity so that we may examine at what ages depression is likely to change, and what may be done at this age to help treat this disease. Given that individuals with higher starting points (intercepts), had steeper trajectories (slopes), it is important to identify depressive symptoms early and when they are increasing as our results suggest that those who start with higher depressive symptoms are at a greater risk of continuing to have higher trajectories of depressive symptoms. These findings have implications for clinical services, schools and parents, who should be made aware that as well as being more likely to experience depression, females are also likely to be younger when depressive symptoms are increasing most rapidly.

Our findings also suggested that the age of maximum depressive symptoms was approximately a year apart between males (20.7 years old) and females (19.7 years old), although the evidence for this was weak. Other research has observed the age of peak depressive symptoms, and similar results are found in studies that use population trajectories with rich longitudinal data. For example Hankin et al., [43] examined depression on 11 to 21 years old and found similar peaks for males and females occurring between the ages of 18 and 21. Sutin et al., [24] found that depressive symptoms were highest at ages 19 and 20 and then decreased, although their research focussed on depressive symptoms from 19 onwards. Natsuaki et al., [36] used multilevel growth-curve models to estimate trajectories of depressive symptoms and their risk factors. Whilst the authors found that females had steeper trajectories compared to males, both sexes peaked in their depressive symptoms at around age 17, several years before the peak in our current study. One possible explanation for this variation in age is the difference in sample. Natsuaki et al., used data from the National Longitudinal Study of Adolescents Health (Add Health), an on-going study that used data from three collection points, with initial study recruitment varying from 12 to 19 years old. One potential problem is that when the waves were assessed, the varying ages may not have had the same characteristics (e.g., assessing depressive symptoms for a 19 year old and a 12 year old concurrently). In contrast, we used data from ALSPAC where the age variation is a maximum of 1 year 8 months, so this variability is minimised. Additionally, we were able to draw measurements from up to eight timepoints, resulting in an increased ability to detect subtle changes in depressive symptoms, rather than more obvious changes from fewer occasions. These contrasting datasets and the number of measurements may help explain the differences in the predicted ages of maximum depressive symptoms. However, what is still not clear is why females and males have differing trajectories of depressive symptoms and why they differ in their ages of peak velocity of depressive symptoms and maximum depressive symptoms.

Thapar et al., [2] suggested that females have higher depressive symptoms in adolescence due to pubertal changes that might result in increased responsiveness to stressors in females. One possible reason why we observed an earlier age of peak velocity of depressive symptoms in females may be through the role of puberty. Women tend to experience puberty earlier than men and evidence has suggested that pubertal status and/or the timing of puberty may be mechanisms responsible for depression [44-46]. For instance, Angold et al., [46] found that depression was higher for females between the ages of 9 and 16, and this seemed to coincide with both pubertal status and timing of puberty. Of note, the number of girls with depression in their study was highest at around age 14, which coincides with the age of peak velocity of depressive symptoms finding in our study, suggesting there may be some mechanism at this age that is underpinning depression and depressive symptoms, and how this manifests.

Research has also shown that an earlier age of menarche is positively associated with higher depressive symptoms [47, 48] and a causal mechanism for depressive symptoms [49]. Transitioning through puberty is also associated with other psychological and social changes, and individuals who transition early may not have developed the cognitive and emotional skills to combat these changes, and therefore experience lasting effects of depressive symptoms. The fact that puberty occurs earlier in women may explain why women have higher trajectories. Our findings suggested that individuals with higher starting points, had higher trajectories. Therefore an earlier age of higher depressive symptoms may set an individual up for a higher trajectory which takes longer to recover from. This could explain why females have higher trajectories compared to males, although more research using the timing and changes in pubertal status for both males and females would be needed to substantiate this claim.

This study has a number of strengths that include using rich longitudinal data that spans over a decade and captures development from late childhood through to young adulthood. Depressive symptoms were measured regularly (at least every 3 years), which allowed to us examine changes in depressive symptoms in greater detail than previous studies. Data with sex and at least one measurement of the SMFQ were available for 9301 individuals, making this one of the larger longitudinal studies on trajectories of depressive symptoms. Using ALSPAC data, it would also be possible to use this model to explore other known predictors of depressive symptoms, for example parity, maternal depression or social deprivation.

A limitation that arises with the data used here, and more generally with longitudinal data, is attrition and the role this plays in biasing results towards individuals who respond. Our sample size decreased from 7335 at the first wave of data collection to 3840 by the eighth occasion opening the possibility to potential bias. Previous studies have imputed missing data in longitudinal studies utilising a missing at random approach (MAR) and found that this bias in ALSPAC is not substantial [10, 50]. The multilevel growth-curves models sued in the present study also assume the missing depressive symptom scores are MAR, so the likelihood of bias is minimalised here.

Additionally, modelling trajectories of depressive symptoms appropriately is challenging and whilst we empirically compared our cubic polynomial model to quadratic and quartic models, other models could in theory be used to measure trajectories of depressive symptoms with potentially better effect. We chose a cubic polynomial model given it is a more parsimonious approach in comparison to splines and fractional polynomials and that it has been used in previous other studies [51-53]. However, a restricted cubic spline approach at certain stages of development may fit the data better. Additionally, cubic terms (and other polynomials) may perform poorly at the start and end of the trajectories, as well as potentially producing artificial turns in the data that do not exist [19]. We checked to ensure that no artificial turns occurred in our data by comparing against other models and the underlying data and descriptive statistics for plausibility. Additionally, the age of peak velocity of depressive symptoms and age of maximum depressive symptoms were calculated well within the trajectories so any bias from potentially mismodelling the start and end of the trajectories is minimised.

In conclusion, we have estimated trajectories of depressive symptoms between males and females in a UK based population cohort across a wide time period. Using multilevel growth-curve models, our findings suggested that females were associated with higher and steeper trajectories of depressive symptoms compared to males, indicating they were more likely to experience higher depressive symptoms for longer. Importantly, the age at which depressive symptoms were increasing most rapidly, and the age of maximum depressive symptoms were much earlier for females compared to males. Whilst the mechanisms underpinning this sex difference are not entirely understood, pubertal status and the timing of pubertal status are likely to play roles in explaining why females have higher trajectories and commence on these trajectories earlier than males. Calculating the age of peak velocity of depressive symptoms is a potentially useful tool for exploring how depressive symptoms are changing and at what point they are increasing most rapidly, which may have consequences downstream. If this can be used for clinical purposes, it may be possible to treat individuals at this age, which may help reduce depressive symptoms or depression at a later stage.

## Acknowledgements

We are extremely grateful to all the families who took part in this study, the midwives for their help in recruiting them, and the whole ALSPAC team, which includes interviewers, computer and laboratory technicians, clerical workers, research scientists, volunteers, managers, receptionists and nurses. This publication is the work of the authors and A.S.F.K and G.L. will serve as guarantors for the contents of this paper. The UK Medical Research Council and Wellcome (Grant Ref: 102215/2/13/2) and the University of Bristol provide core support for ALSPAC. A comprehensive list of grants funding is available on the ALSPAC website. This research was specifically funded by Wellcome (08426812/Z/07/Z), Wellcome and the MRC (076467/Z/05/Z; 092731; 092731; 092731), the MRC (MR/M006727/1), NIH (PD301198-SC101645). A.S.F.K is funded by an ESRC Advanced Quantitative Methods Studentship. NJT is a Wellcome Trust Investigator (202802/Z/16/Z), is a programme lead in the MRC Integrative Epidemiology Unit (MC_UU_12013/3) and works within the University of Bristol NIHR Biomedical Research Centre (BRC) and CRUK Integrative Cancer Epidemiology Programme (C18281/A19169).

## Supplementary Materials

Supplementary materials and example Stata code can be found here: XXXXXX

## Author Contributions

A.S.F.K, D.M, N.J.T, R.P, J.H and G.L conceived the study design. A.S.F.K performed the analysis under the supervision of G.L. A.S.F.K wrote the initial draft of the manuscript. All authors contributed to and approved the final manuscript.

